# Interpretable Machine Learning Links Group-Specific Plasma Proteomics to Regional Structural Neuroimaging in Alzheimer’s Disease

**DOI:** 10.64898/2026.11.02.763368

**Authors:** Hemshankar Laugi, Mohammadreza Hosseinzadehketilateh, Seonjoo Lee, Steven M. Weisberg, Xi Zhu, the Alzheimer’s Disease Neuroimaging Initiative

## Abstract

**Background:** Alzheimer’s disease (AD) pathology begins years before clinical symptoms, highlighting the need for scalable biomarkers that capture biological changes across the AD continuum. Broader plasma proteomics may capture systemic processes that vary with disease stage, yet their relationship to AD-related brain neurodegeneration remains unclear. We aimed to identify plasma proteomic signatures across cognitively normal (CN), mild cognitive impairment (MCI), and AD and evaluate their neuroanatomical associations.

**Methods:** We analyzed 146 plasma analytes from 565 Alzheimer’s Disease Neuroimaging Initiative (ADNI) participants (58 CN, 396 MCI, 111 AD). Group-specific proteomic signatures were identified using pairwise classification models with nested 5×5 cross-validation. SHAP quantified feature importance, with the top 15 proteins per comparison being prioritized for MRI association analysis. We analyzed 103 regional structural MRI volumes harmonized across 57 acquisition sites using ComBat. Separate ordinary least squares regression tested protein-MRI associations, adjusting for age, sex, APOE ε4, diagnosis, and intracranial volume, followed by Benjamini-Hochberg false discovery rate (FDR) correction.

**Results:** Classification models using proteomic data achieved AUC of 0.956 ± 0.017 for CN vs MCI, 0.963 ± 0.011 for MCI vs. AD, and 0.836 ± 0.085 for CN vs. AD. SHAP identified group-dependent proteomic signatures. No protein-MRI associations survived FDR correction for CN vs. MCI. In MCI vs. AD, higher cortisol was associated with lower left precuneus and superior parietal volumes, while higher osteopontin was associated with lower left paracentral and accumbens volumes. In CN vs. AD, higher vitronectin was associated with lower posterior corpus callosum volume.

**Conclusion:** These findings suggest that plasma proteomics capture group-specific systemic biological signals across the AD continuum. Linking ML-prioritized plasma proteins to regional brain structure provides neuroanatomical context for blood-based biomarker discovery.

## 1. Introduction

Alzheimer’s disease (AD) is a progressive neurodegenerative disorder characterized by gradual cognitive decline and molecular, cellular, and structural brain changes. Accurate characterization of AD across clinically defined groups, including cognitively normal (CN), mild cognitive impairment (MCI), and AD, is important for earlier detection, better biological understanding, and development of targeted intervention strategies. In routine practice, initial evaluation still relies heavily on cognitive testing, questionnaires, and clinical assessment. However, these approaches do not directly measure the biological processes driving the disease. Hence, there is an increasing need to incorporate biomarkers alongside cognitive symptoms to provide a biologically grounded definition of AD and to link cognitive decline to underlying disease pathology (Dubois et al., 2014; Hansson et al., 2022; Jack et al., 2018, 2024).

The progression of AD begins years before clinical symptoms become apparent, with amyloid-β (Aβ) and tau pathology generally preceding detectable neurodegeneration (Jack et al., 2018, 2024; Krix et al., 2024). Existing diagnostic tools, such as cerebrospinal fluid (CSF) biomarkers, positron emission tomography (PET), and magnetic resonance imaging (MRI), provide complementary information about these underlying disease processes. CSF and PET characterize molecular pathology related to amyloid and tau, whereas structural MRI provides noninvasive measures of neurodegeneration through changes in regional brain volume and cortical thickness (Blennow & Zetterberg, 2018; Frisoni et al., 2010; Jack et al., 2018, 2024; Johnson et al., 2012). However, CSF collection is invasive, PET is expensive and has limited accessibility, and repeated advanced neuroimaging is not practical for population-level screening. Hence, there is an urgent need for scalable, minimally invasive, and biologically informative biomarkers for early detection, screening, and monitoring of AD-related change (Frisoni et al., 2025; Palmqvist et al., 2025).

Blood-based biomarkers have emerged as a promising and scalable approach for addressing these limitations. Large-scale plasma proteomic profiling can capture a broad range of disease-associated biological processes, including immune, vascular, metabolic, lipid, and neurodegeneration-related pathways, providing information beyond a small number of established AD biomarkers (Heo et al., 2025). However, circulating proteins reflect both systemic and central nervous system-related processes and may be influenced by inflammation, vascular function, lipid metabolism, and other physiological processes that do not necessarily correspond to brain structural abnormalities (Heo et al., 2025; Morgan et al., 2019; Sweeney et al., 2018). Although previous studies have used plasma proteomics to distinguish clinical groups (Heo et al., 2025; Llano et al., 2013), proteins contributing to classification may primarily reflect systemic physiological differences without necessarily showing measurable associations with regional brain structure.

Structural MRI is particularly useful for investigating this relationship because AD-related neurodegeneration follows regionally heterogeneous patterns rather than affecting the brain uniformly. Regional structural alterations have been reported across cortical and subcortical regions affected by AD, making MRI a useful complementary modality for evaluating whether peripheral biological signals correspond to regional brain structure (Dickerson et al., 2009). Previous studies have linked circulating proteins with structural brain measures, including associations between plasma clusterin and longitudinal brain atrophy in MCI (Thambisetty et al., 2011), and a recent scoping review identified 63 studies examining peripheral proteins in relation to MRI-derived measures in the context of AD research, with grey matter volume being the most frequently examined imaging measure and inflammatory and immune-related proteins representing many of the reported associations (Li et al., 2026). Conversely, studies focusing on protein-MRI associations can identify circulating proteins related to brain structure, but these associations alone do not establish whether those proteins also contribute to distinguishing clinically defined CN, MCI, and AD groups.

The primary aim of this study was to determine whether plasma proteins that contribute to distinguishing clinical groups also show regional neuroanatomical associations, providing biological context beyond classification performance alone. We hypothesized that a subset of group-discriminative plasma proteins would also show associations with regional brain volumes. We integrated interpretable machine learning with regional structural neuroimaging to examine plasma proteomic signatures across three clinically defined comparisons: CN vs. MCI, MCI vs. AD, and CN vs. AD. Using data from the Alzheimer’s Disease Neuroimaging Initiative (ADNI), we applied machine learning (ML) analysis to identify group-specific plasma proteins and carried the top-ranked proteins from each comparison forward for MRI integration. We then tested whether these identified proteins were associated with regional cortical and subcortical volumes.

## Methods

### 2.1 Study population and data sources

We used baseline data from the Alzheimer’s Disease Neuroimaging Initiative (ADNI), a longitudinal, multicenter public dataset focused on AD biomarkers (Mueller et al., 2005). This dataset was accessed upon registration and approval in accordance with the data use agreements. The diagnostic criteria in ADNI are based on a composite clinical assessment that includes measures such as the Mini-Mental State Examination (MMSE), the Clinical Dementia Rating (CDR), and the Logical Memory tests. Participants were categorized into three clinical groups: cognitively normal (CN), mild cognitive impairment (MCI), and Alzheimer’s disease (AD), with early MCI (EMCI) and late MCI (LMCI) combined into a single MCI group. We used ADNI-provided clinical diagnoses, which distinguish CN participants without significant cognitive impairment (CDR = 0), MCI participants with memory impairment but largely preserved daily functioning (CDR = 0.5), and AD participants with clinically diagnosed dementia (CDR generally 0.5 or 1), together with MMSE, Logical Memory performance, and clinician judgment.

The initial cohort included 566 subjects with plasma proteomics and matched structural MRI data (58 CN, 396 MCI, and 112 AD). Each subject had 146 protein analytes measured via the Luminex xMAP platform. The final cohort used for this research study was also screened for flagged participants, and no participants associated with known data-quality concerns were identified. Group differences were evaluated using one-way analysis of variance (ANOVA) for the continuous variable (age) and chi-square tests for categorical variables (sex and APOE ε4 carrier status).

### 2.2 Proteomic data preprocessing and covariate adjustment

During initial preprocessing, no missing values were reported for any of the 146 plasma analytes among 566 subjects. To remove demographic and genetic effects, each protein feature was adjusted for age, sex, and APOE ε4 status via linear regression applied within training folds only. This residualization step was performed so that the ML and SHAP analyses were more likely to reflect proteomic signals associated with disease progression rather than demographic or genetic effects. After residualization, feature normalization was applied based on each feature’s skewness. Z-score normalization was applied for features with low skewness (|skewness| ≤ 1), whereas the Yeo-Johnson transformation was performed for features with higher skewness (|skewness| > 1). These preprocessing steps were done within training folds only to prevent data leakage and ensure unbiased evaluation. Hence, the final analysis included 566 subjects, each with 146 covariate-adjusted and normalized plasma protein biomarkers.

### 2.3 Machine learning framework and evaluation on proteomic data

Five supervised machine learning models were applied: Elastic Net logistic regression, support vector machine (SVM) with radial basis function (RBF) kernel, random forest, XGBoost, and naïve Bayes across three clinical group comparisons (CN vs MCI, MCI vs AD, and CN vs AD). A nested stratified 5×5 cross-validation framework was applied to evaluate model performance and ensure unbiased generalization. As shown in Figure 1, the outer loop was partitioned into 5 stratified folds, where in each iteration, 80% of the data was used for training and validation, and 20% was held out as an independent test fold for unbiased performance estimation. The training sets from each outer split were used in the inner loop, which again consisted of 5 stratified folds, with each fold serving as a validation set once. For hyperparameter tuning, grid search was performed with cross-validation across the inner folds. Model selection within the inner loop was based on mean validation AUC across the inner folds. The optimized model from the inner loop was then retrained on the full outer training set and evaluated on the corresponding outer test fold. This process was repeated across all outer folds.

**Figure 1.**
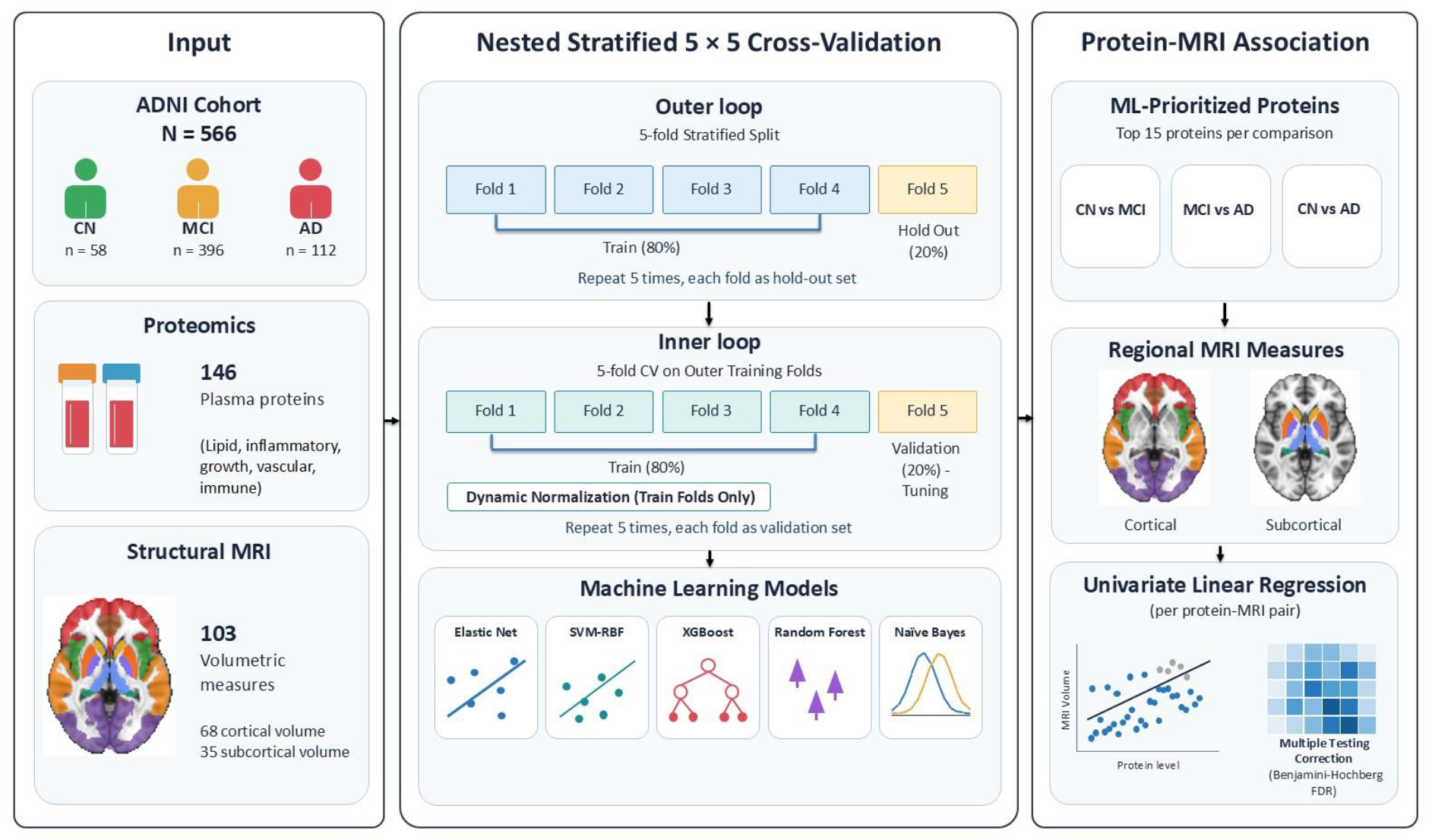
Integrated machine learning and protein-MRI association framework.

The model performance was evaluated using the Receiver Operating Characteristic -Area Under the Curve (ROC AUC) as the primary metric. Accuracy, sensitivity, and specificity of each model across each task were also reported as secondary metrics. Each performance metric was averaged across outer folds and reported as mean ± standard deviation. Class imbalance was handled using model-specific weighting strategies, including class weighting in linear and tree-based models and scale_pos_weight in XGBoost.

### 2.4 Model Interpretability via Out-of-Sample SHAP Analysis

SHapley Additive exPlanations (SHAP) were performed to quantify the contribution of individual plasma proteins to model predictions and identify key features across classification tasks. The XGBoost model was used because of their strong predictive performance and SHAP compatibility. To ensure out-of-sample interpretability, SHAP values were computed only on outer test folds within the nested cross-validation framework for each classification task.

SHAP analysis resulted in two outputs: global feature importance (bar plots) and feature directionality (beeswarm plots). Mean absolute SHAP values were used to rank proteins by their overall contribution to model predictions, while beeswarm plots were used to visualize the direction of feature contributions. Top-ranking proteins were further grouped into broader functional categories to characterize group-specific proteomic patterns. For MRI integration, the 15 highest-ranked proteins from each comparison were selected as a focused and consistent subset of the 146-analyte panel, balancing retention of highly influential SHAP features with the multiple-testing burden of subsequent regional MRI association analyses.

### 2.5 Structural MRI Integration and Protein-MRI Association Analysis

Regional structural MRI measures were obtained from the ADNI/UCSF FreeSurfer 7.2.0 processed dataset. FreeSurfer provides automated cortical parcellation and segmentation of structural MRI, allowing regional cortical and subcortical morphometric measures to be quantified (Fischl, 2012; Fischl et al., 2002). Cortical regions were based on the Desikan-Killiany atlas, which subdivides the cerebral cortex into anatomically defined gyral-based regions of interest (Desikan et al., 2006).

We matched the FreeSurfer dataset to the baseline clinical and proteomics cohort using subject IDs. For each participant, the MRI scan closest to the baseline clinical/proteomics examination date was selected. MRI data were available for all 566 subjects included in the proteomics cohort, with a median absolute interval of 16 days and a mean interval of 21.2 days between the selected MRI scan and baseline examination.

The final MRI feature set consisted of 103 regional volumes, including 68 cortical volumes and 35 subcortical volumes. Intracranial volume (ICV) was retained as a covariate rather than an MRI outcome.

Because MRI data were acquired across multiple ADNI sites, regional MRI measures were harmonized using ComBat to reduce site-related variability while preserving biological variation associated with age, sex, APOE ε4 status, diagnosis, and ICV. One participant representing the only observation from an MRI acquisition site was excluded before harmonization, resulting in a final MRI association cohort of 565 participants (58 CN, 396 MCI, and 111 AD) across 57 acquisition sites.

Protein-MRI association analyses were conducted separately for CN vs. MCI (N=454), MCI vs. AD (N=507), and CN vs. AD (N=169). For each comparison, the 15 plasma proteins prioritized by SHAP were individually tested against each of the 103 regional volumes using separate univariate ordinary least-squares (OLS) regression models. Each model included one plasma protein as the predictor of interest and one regional MRI volume as the outcome, adjusting for age, sex, APOE ε4 status, diagnosis, and ICV.

This resulted in 1,545 protein-MRI association tests within each clinical-group comparison (15 proteins × 103 MRI measures). Benjamini-Hochberg false discovery rate (FDR) correction was applied separately across the 1,545 tests within each comparison, with FDR-adjusted p < 0.05 considered statistically significant.

## 2. Results

### 3.1 Cohort characteristics

The three clinical groups did not differ significantly in age (CN: 75.1 ± 5.8 years; MCI: 74.7 ± 7.4 years; AD: 74.8 ± 8.1 years; p = 0.939) or sex distribution (p = 0.104). APOE ε4 carrier status differed significantly across groups (p < 0.001), with carriers comprising 8.6% (5/58) of CN, 53.3% (211/396) of MCI, and 67.9% (76/112) of AD participants.

### 3.2 Classification performance of proteomic data

We evaluated model performance across three clinical-group comparisons: CN vs. MCI, MCI vs. AD, and CN vs. AD. The performance of the best model in each comparison is summarized in Table 1. SVM (RBF) performed best, with outer test AUCs of 0.956 ± 0.017 in CN vs MCI and 0.963 ± 0.011 in MCI vs AD. However, random forest performed best on the CN vs AD task, with an outer test AUC of 0.836 ± 0.085.

**Table 1.** Model performance across classification tasks. Performance of the best-performing model for each classification task using covariate-adjusted plasma proteomic features. Values are reported as mean ± standard deviation across outer cross-validation folds.

| **Classification Task** | **Best Model** | **AUC (mean ± SD)** | **Accuracy** | **Sensitivity** | **Specificity** |
|---|---|---|---|---|---|
| **CN vs MCI** | SVM (RBF) | 0.956 ± 0.017 | 0.918 ± 0.013 | 1.000 ± 0.000 | 0.362 ± 0.099 |
| **MCI vs AD** | SVM (RBF) | 0.963 ± 0.011 | 0.905 ± 0.014 | 0.730 ± 0.118 | 0.954 ± 0.027 |
| **CN vs AD** | Random Forest | 0.836 ± 0.085 | 0.759 ± 0.066 | 0.982 ± 0.022 | 0.327 ± 0.180 |

**Table 2:** Significant associations between SHAP-prioritized plasma proteins and regional MRI volumes. β represents the regression coefficient and SE its standard error; negative β values indicate associations between higher protein levels and lower regional volume.

| **Comparison** | **Plasma protein** | **MRI region** | **N** | **β** | **SE** | ***p*-value** | **FDR-adjusted *q*-value** |
|---|---|---|---|---|---|---|---|
| MCI vs. AD | Cortisol | Left precuneus | 507 | −1213.84 | 282.96 | 2.1 × 10<sup>−5</sup> | 0.0285 |
| MCI vs. AD | Cortisol | Left superior parietal | 507 | −1802.09 | 439.73 | 4.9 × 10<sup>−5</sup> | 0.0285 |
| MCI vs. AD | Osteopontin | Left paracentral | 507 | −304.49 | 74.87 | 5.5 × 10<sup>−5</sup> | 0.0285 |
| MCI vs. AD | Osteopontin | Left accumbens area | 507 | −61.65 | 15.46 | 7.6 × 10<sup>−5</sup> | 0.0295 |
| CN vs. AD | Vitronectin | Posterior corpus callosum | 169 | −711.39 | 161.06 | 1.8 × 10<sup>−5</sup> | 0.0282 |

### 3.3 SHAP-based identification of group-specific plasma proteins

SHAP analysis was used to understand the individual contribution of each proteomic biomarker to model performance (Figure 2). In the early stage (CN vs MCI) category, the most important features were associated with lipid metabolism (e.g., apolipoprotein A-II and apolipoprotein D), transport (e.g., transthyretin and serotransferrin), and early inflammatory or immune activity (e.g., Interleukin-16). The late-stage (MCI vs AD) task showed a shift towards growth-related, inflammatory, and complement-associated signaling proteins as the top predictors. PAPP-A (Pregnancy-Associated Plasma Protein A), which regulates IGF (insulin-like growth factor) signaling, emerged as the leading feature. Other important proteins include growth factor–related proteins such as Heparin-binding Epidermal Growth Factor (HB-EGF) and insulin-like growth factor binding proteins (IGFBPs), and complement pathway components such as Complement Factor H. In the CN vs AD classification task, some of the most important proteins included Serum Glutamic Oxaloacetic Transaminase (SGOT), Betacellulin, Brain Natriuretic Peptide, Matrix Metalloproteinase-2 (MMP-2), and Tenascin-C. SHAP directionality (beeswarm plots) further helped to understand whether higher or lower protein values pushed predictions toward one disease class or the other. Many top features showed consistent directional trends across folds, which implies that the identified signals were stable and not driven by a few outlier samples. The top 15 SHAP-ranked proteins from each classification task were subsequently evaluated for associations with regional structural MRI measures.

**Figure 2.**
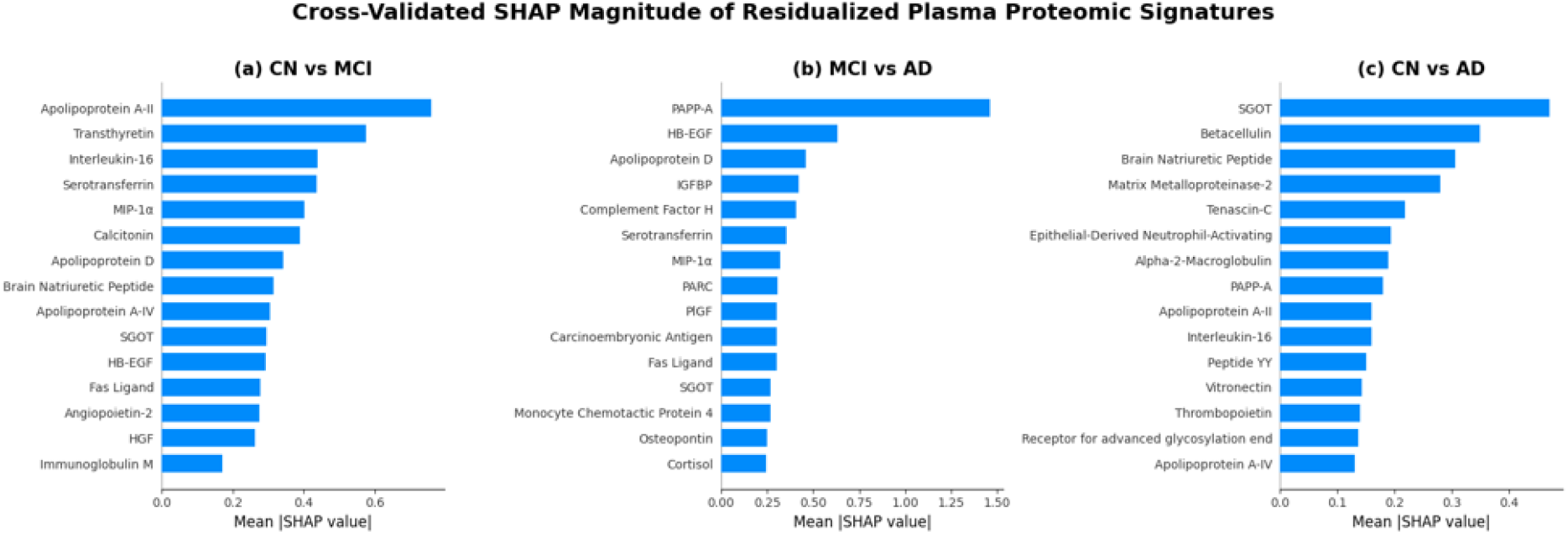
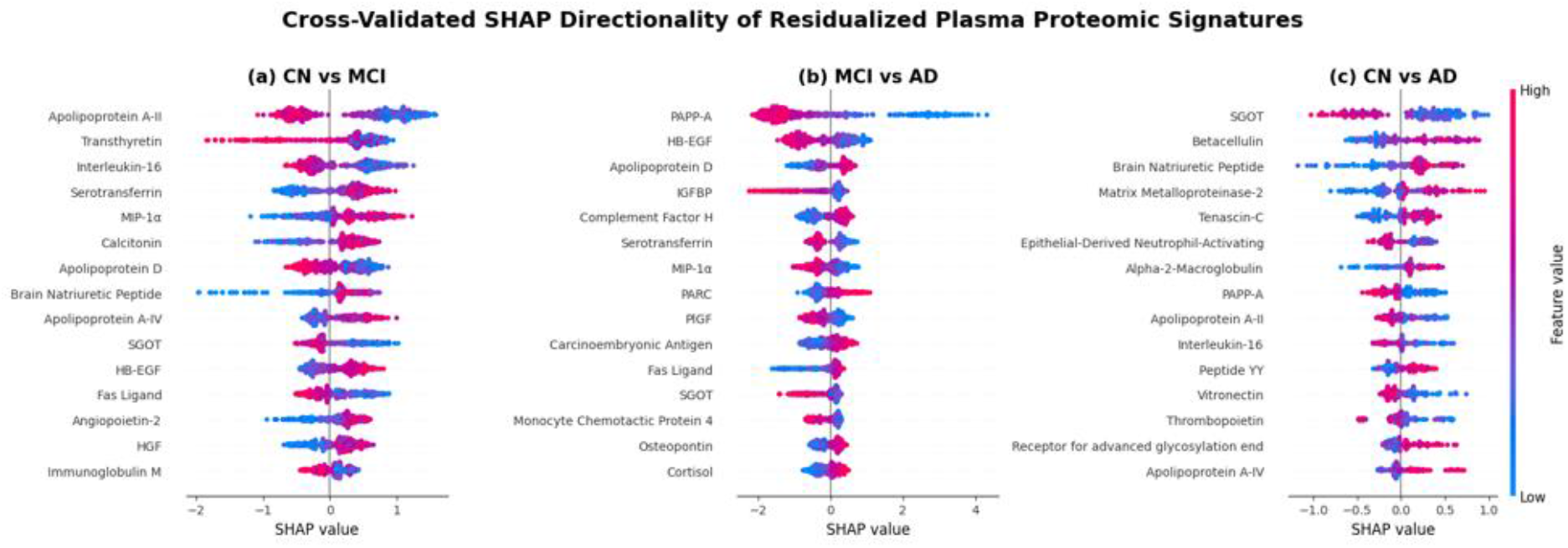
Stage-specific feature importance identified using SHAP. 2 (i) Global feature importance (bar plots) and 2 (ii) feature directionality (beeswarm plots) for each classification task based on XGBoost models. SHAP values were computed on held-out outer folds and aggregated across folds to reflect out-of-sample feature contributions.

**Figure 3:**
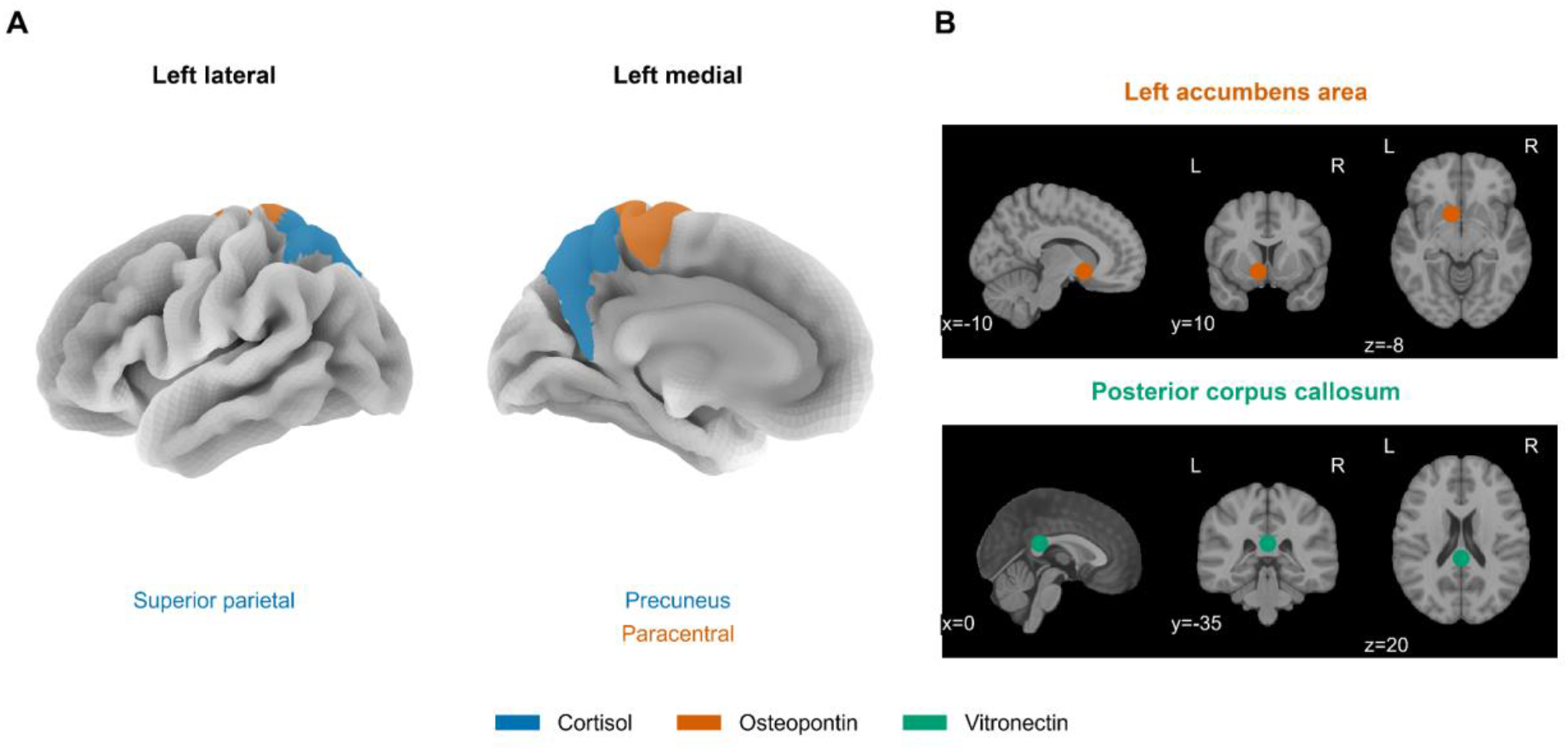
Anatomical distribution of significant plasma protein-MRI associations. (A) Cortical regions associated with cortisol (blue) and osteopontin (orange) in the MCI vs. AD comparison. (B) Approximate locations of the left accumbens area associated with osteopontin (orange) and posterior corpus callosum associated with vitronectin (green); all displayed associations survived FDR correction (*q* < 0.05), and markers in panel B indicate anatomical locations rather than voxelwise statistical results.

### 3.4 Associations Between SHAP-Prioritized Plasma Proteins and Regional Brain Structure

The CN vs. MCI analysis included 454 subjects (58 CN and 396 MCI). A total of 114 protein-MRI associations had nominal p < 0.05; however, none survived FDR correction (minimum q = 0.299). The MCI vs. AD analysis included 507 subjects (396 MCI and 111 AD). A total of 137 associations had nominal p < 0.05, of which four remained significant after FDR correction. All four associations had negative regression coefficients. Higher cortisol was associated with lower left precuneus volume (β = −1213.84, q = 0.0285) and lower left superior parietal volume (β = −1802.09, q = 0.0285). Higher osteopontin was associated with lower left paracentral volume (β = −304.49, q = 0.0285) and lower left accumbens area volume (β = −61.65, q = 0.0295). The CN vs. AD analysis included 169 subjects (58 CN and 111 AD). A total of 58 associations had nominal p < 0.05, of which one remained significant after FDR correction. Higher vitronectin was associated with lower posterior corpus callosum volume (β = −711.39, q = 0.0282).

## Discussion

This study identified distinct group-discriminative plasma proteomic signatures using interpretable machine learning, but only five protein-MRI associations survived FDR correction. The CN vs. MCI comparison highlighted lipid-transport and immune-related proteins without significant regional MRI associations. In MCI vs. AD, cortisol and osteopontin were associated with lower volumes in four brain regions, while in CN vs. AD, vitronectin was associated with lower posterior corpus callosum volume. Classification performance was higher for CN vs. MCI and MCI vs. AD than for CN vs. AD. These findings suggest that proteins contributing to clinical-group discrimination do not necessarily show measurable relationships with regional brain structure.

In CN vs. MCI, several highly ranked proteins were involved in lipid metabolism and transport, including apolipoproteins A-II, A-IV, and D, and transthyretin, together with immune-related proteins such as IL-16 and MIP-1α. This pattern is biologically plausible because disturbances in lipid homeostasis, particularly those involving apolipoprotein pathways, are closely connected to AD biology. APOE has major roles in lipid transport and neuronal homeostasis, while APOE ε4 influences multiple processes implicated in AD, including lipid metabolism, neuroinflammation, and vascular function (Liu et al., 2013). Our findings are broadly consistent with recent large-scale plasma proteomic studies identifying AD-associated proteins involved in lipid dysregulation and immune responses (Heo et al., 2025). Therefore, the combination of lipid/transport and immune proteins may reflect systemic processes that differentiate CN and MCI.

The proteomic pattern was different in MCI vs. AD, with proteins related to inflammatory, complement, growth-factor, and vascular processes, including PAPP-A, HB-EGF, IGFBP, complement factor H, MIP-1α, PARC, PlGF, and osteopontin. This pattern is consistent with the increasingly recognized role of sustained immune and inflammatory responses in AD. Complement pathways can contribute to both normal homeostatic functions and inflammatory and synaptic processes relevant to neurodegeneration, while microglial, astrocytic, and vascular responses form part of the complex cellular response to AD pathology (De Strooper & Karran, 2016; Tenner et al., 2018). For CN vs. AD, the broader mixture of metabolic, vascular, inflammatory, extracellular-matrix, and neuroendocrine-related proteins may similarly reflect the multisystem nature of the clinical differences between these groups. Importantly, Llano et al. (2013), who analyzed the same 146-analyte ADNI plasma panel, identified apolipoprotein A-II, SGOT, and BNP as recurring proteins across their AD-versus-control classification signatures; all three were also among our top 15 SHAP-ranked proteins in CN vs. AD. Although our broader lipid and immune-related findings align with previous studies, the specific proteins and their relative importance differ across studies, potentially reflecting differences in study populations, diagnostic comparisons, proteomic platforms, and analytical methods (Heo et al., 2025; Llano et al., 2013). Importantly, these patterns should be viewed as group-specific signatures rather than a sequential biological progression from one pathway to another.

Cortisol was among the 15 SHAP-prioritized proteins in MCI vs. AD and showed negative associations with left precuneus and superior parietal volumes. Cortisol is a glucocorticoid regulated by the hypothalamic-pituitary-adrenal (HPA) axis, and chronic HPA-axis dysregulation and prolonged glucocorticoid exposure have been linked to cognitive impairment and neurodegenerative processes (Ouanes & Popp, 2019). In an ADNI study of 304 individuals with amnestic MCI, higher baseline plasma cortisol predicted a faster decline in hippocampal volume over time (White et al., 2023). Our findings are consistent with a broader association between circulating cortisol and brain structure, although the affected regions differ from the hippocampal finding reported by White et al., potentially reflecting differences in study design and regional MRI outcomes. The precuneus is particularly relevant because structural MRI studies have demonstrated prominent precuneus atrophy in AD, including early-onset AD (Karas et al., 2007).

Osteopontin was also prioritized in MCI vs. AD and was associated with lower left paracentral and accumbens volumes. Osteopontin is an inflammatory mediator that has been investigated in both AD and vascular cognitive impairment, providing a potential link between immune/vascular processes and neurodegeneration. Chai et al. (2021) reported higher plasma osteopontin across cognitive-impairment groups and demonstrated relationships with inflammatory markers, cerebrovascular disease, and neuroimaging measures of brain atrophy. Rather than implying that osteopontin specifically targets these regions, a more plausible interpretation is that circulating osteopontin may index an inflammatory and vascular environment associated with structural brain vulnerability.

The vitronectin finding introduces a different biological pathway. Vitronectin is an abundant multifunctional glycoprotein found in plasma and the extracellular matrix and has established roles in cell adhesion, tissue remodeling, hemostatic and vascular processes, and regulation of the complement system (Preissner & Reuning, 2011; Schvartz et al., 1999). Its relevance to AD is supported by earlier neuropathological evidence showing vitronectin immunoreactivity in senile plaques and neurofibrillary tangles in AD brain tissue (Akiyama et al., 1991). More recent evidence further strengthens the connection with AD-related amyloid biology. A 2026 study of the plasma albumin interactome identified vitronectin as a key hub protein that was reduced in amyloid-PET-positive individuals; functional experiments further showed that vitronectin directly bound Aβ and inhibited its oligomerization (Kang et al., 2026). Therefore, the association between plasma vitronectin and posterior corpus callosum volume may reflect a relationship between systemic extracellular-matrix, vascular, or complement-related biology and structural brain integrity; its recently reported relationship with plasma Aβ biology provides an additional potential connection to AD. However, unlike cortisol and osteopontin, there is currently limited evidence directly linking circulating vitronectin to regional MRI volume, and particularly to the posterior corpus callosum. The regional association should therefore be interpreted as a novel hypothesis-generating finding rather than evidence for an established vitronectin-mediated mechanism.

The absence of significant MRI associations in CN vs. MCI is also informative when considered alongside the SHAP results. Although the CN vs. MCI models identified strong group-discriminative proteomic signals, particularly involving lipid transport and immune-related proteins, these proteins did not show sufficiently strong relationships with regional brain volumes after multiple-testing correction. This may indicate that some circulating signals reflect systemic biology not strongly coupled to macroscopic regional volume, while other structural relationships may be smaller, spatially distributed, or better captured by different imaging phenotypes. The smaller CN sample may also have reduced statistical power. Therefore, the absence of FDR-supported associations should not be interpreted as evidence that these proteins lack biological relevance to the brain.

Several limitations should be considered when interpreting these results. First, the current analysis relies on cross-sectional data, which limits the ability to assess long-term disease progression or establish causal relationships between circulating proteins and structural brain differences. CN, MCI, and AD are also clinically defined groups and do not perfectly represent underlying biological AD status. Therefore, the group-specific proteomic patterns identified here should not be interpreted as direct temporal stages of AD pathology. Future analyses incorporating amyloid and tau biomarkers would help determine whether the identified proteomic and MRI relationships are specifically associated with biological AD pathology beyond clinical diagnosis. Second, the CN group was relatively small compared with the MCI group, which may have increased uncertainty in comparisons involving CN and reduced power for the MRI analyses. The relatively lower and more variable CN vs. AD classification performance (AUC = 0.836 ± 0.085) should also be interpreted cautiously. Although the smaller comparison cohort (N = 170) may have contributed to greater variability, the reason for the lower discrimination remains uncertain. Furthermore, ADNI recruited participants for a research cohort rather than through population-based sampling, which may limit the representativeness of the small CN group (Petersen et al., 2010). Third, MRI and baseline clinical/proteomic assessments were not obtained on exactly the same day for every participant. Although the selected scans were generally close to baseline, this temporal difference may introduce additional variability into the observed protein-MRI relationships.

The protein-MRI analyses were also restricted to the 15 highest-ranked SHAP proteins from each comparison. This provided a consistent and focused subset of the 146-analyte panel while reducing the downstream multiple-testing burden, but potentially relevant relationships involving lower-ranked proteins were not evaluated. There is no universally established SHAP cutoff for multimodal biomarker integration, and future studies should evaluate the robustness of the findings across alternative feature-selection thresholds. Finally, this study does not include independent external validation. Nested cross-validation provides strong internal evaluation of classification performance, while ComBat harmonization and FDR correction address site-related variation and multiple testing, respectively; however, these procedures do not substitute for independent external replication. Future work should therefore validate the cortisol, osteopontin, and vitronectin associations in independent cohorts and use longitudinal plasma and MRI measurements to determine whether changes in these proteins precede, accompany, or follow structural brain changes. Incorporating amyloid and tau biomarkers, additional imaging phenotypes, and independent proteomic platforms may further clarify the biological mechanisms underlying these systemic-to-brain relationships.

## 3. Conclusion

In summary, this study demonstrates the value of integrating interpretable machine learning with structural neuroimaging to move beyond plasma-based classification alone. Broad plasma proteomics captured different systemic biological signatures across CN vs. MCI, MCI vs. AD, and CN vs. AD, including lipid/transport, immune, inflammatory, vascular, growth-factor, extracellular-matrix, and neuroendocrine-related processes. However, MRI integration showed that only a subset of these group-discriminative signals also had measurable relationships with regional brain structure. The structural associations involving cortisol, osteopontin, and vitronectin provide neuroanatomical context for systemic proteomic signals related to neuroendocrine, inflammatory/vascular, and extracellular-matrix-related biological processes.

Overall, these findings highlight an important distinction between proteins that are useful for distinguishing clinical groups and proteins that also show neuroanatomical associations. By using machine learning to prioritize circulating proteins and structural MRI to provide additional biological context, this study provides a focused framework for investigating how systemic biological signals relate to the brain in AD. Longitudinal analysis, biological AD characterization using amyloid and tau biomarkers, and independent replication will be important for determining whether these relationships are reproducible and how they change over time.

## Data Availability

The data used in this study were obtained from the Alzheimer’s Disease Neuroimaging Initiative (ADNI) database (https://adni.loni.usc.edu). Access to the data is available to approved researchers following registration and compliance with the ADNI Data Use Agreement. The authors cannot redistribute individual-level ADNI data under this agreement.

## Acknowledgments

We acknowledge the Alzheimer’s Disease Neuroimaging Initiative (ADNI) and its participating investigators for collecting and providing the clinical, plasma proteomic, and neuroimaging data used in this study. ADNI investigators contributed to the design, implementation, and data collection of ADNI but did not participate in the analysis or writing of this manuscript. A complete list of ADNI investigators is available at https://adni.loni.usc.edu. Data collection and sharing for ADNI is funded by the National Institute on Aging (National Institutes of Health Grant U19 AG024904). The grantee organization is the Northern California Institute for Research and Education. In the past, ADNI has also received funding from the National Institute of Biomedical Imaging and Bioengineering, the Canadian Institutes of Health Research, and private-sector contributions through the Foundation for the National Institutes of Health (FNIH), including contributions from AbbVie; Alzheimer’s Association; Alzheimer’s Drug Discovery Foundation; Araclon Biotech; BioClinica, Inc.; Biogen; Bristol-Myers Squibb Company; CereSpir, Inc.; Cogstate; Eisai Inc.; Elan Pharmaceuticals, Inc.; Eli Lilly and Company; EuroImmun; F. Hoffmann-La Roche Ltd and its affiliated company Genentech, Inc.; Fujirebio; GE Healthcare; IXICO Ltd.; Janssen Alzheimer Immunotherapy Research & Development, LLC; Johnson & Johnson Pharmaceutical Research & Development LLC; Lumosity; Lundbeck; Merck & Co., Inc.; Meso Scale Diagnostics, LLC; NeuroRx Research; Neurotrack Technologies; Novartis Pharmaceuticals Corporation; Pfizer Inc.; Piramal Imaging; Servier; Takeda Pharmaceutical Company; and Transition Therapeutics.

## Notes

### Competing Interest Statement

The authors have declared no competing interest.

