## Supplementary Materials for "Interpretable Machine Learning Links Group-Specific Plasma Proteomics to Regional Structural Neuroimaging in Alzheimer’s Disease"

**Supplementary Figures:**

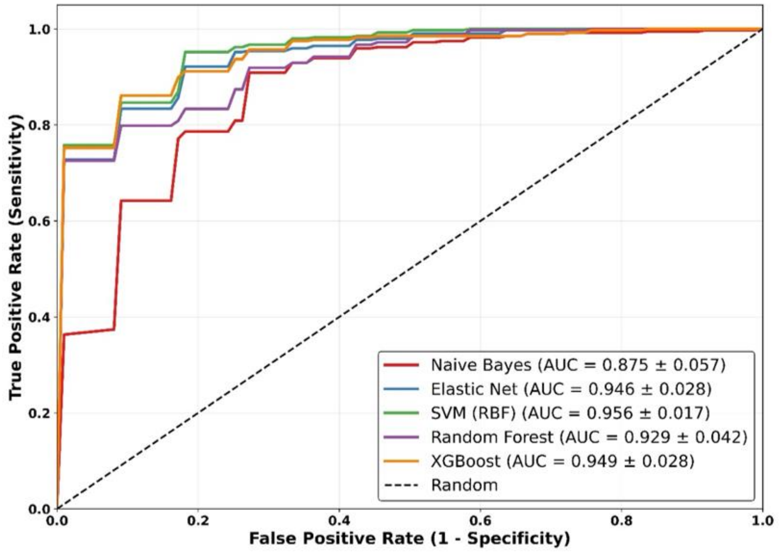


Figure 1 (i): Mean ROC Curve for CN vs MCI

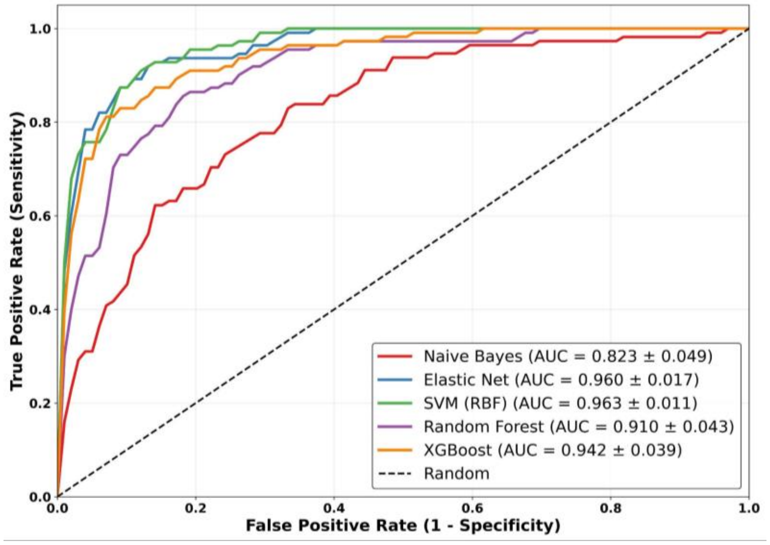


Figure 1 (ii): Mean ROC Curve for MCI vs AD

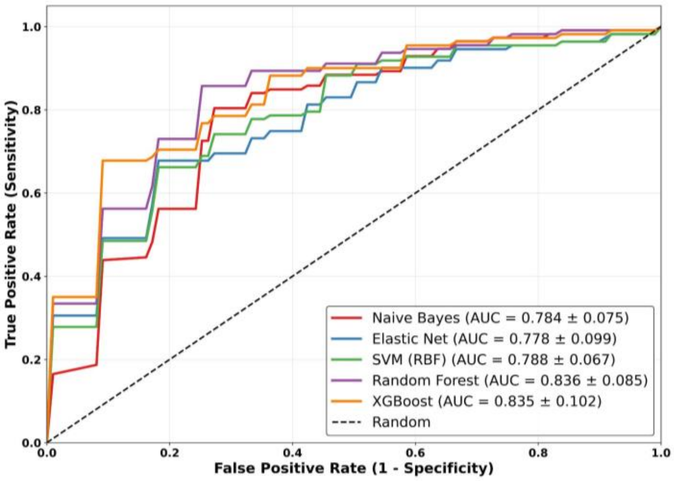


Figure 1 (iii): Mean ROC Curve for CN vs AD

Figure 1. Classification performance across the Alzheimer’s disease continuum

**Table 1: Demographic and Clinical Characteristics of the Study Cohort.**

Continuous variables are reported as mean ± standard deviation (SD), and categorical variables are reported as number (percentage). Group differences were evaluated using one-way analysis of variance (ANOVA) for age and chi-square tests for categorical variables (sex and APOE ε4 carrier status)

| **Characteristics** | **Overall (N=566)** | **CN (N=58)** | **MCI (N=396)** | **AD (N=112)** | **p-value** |
|---|---|---|---|---|---|
| Age, years | 74.8 ± 7.4 | 75.1 ± 5.8 | 74.7 ± 7.4 | 74.8 ± 8.1 | 0.939 |
| Female sex, n (%) | 215 (38.0%) | 28 (48.3%) | 140 (35.4%) | 47 (42.0%) | 0.104 |
| APOE ε4 carrier, n (%) | 292 (51.6%) | 5 (8.6%) | 211 (53.3%) | 76 (67.9%) | <0.001 |
